# EryDB: a transcriptomic profile database for erythropoiesis and erythroid-related diseases

**DOI:** 10.1101/2023.12.22.572961

**Authors:** Guangmin Zheng, Song Wu, Zhaojun Zhang, Zijuan Xin, Lijuan Zhang, Siqi Zhao, Jing Wu, Yanxia Liu, Meng Li, Xiuyan Ruan, Yiming Bao, Hongzhu Qu, Xiangdong Fang

## Abstract

Erythropoiesis is a finely regulated and complex process that involves multiple transformations from hematopoietic stem cells to mature red blood cells at hematopoietic sites from the embryonic to the adult stages. Investigations into its molecular mechanisms have generated a wealth of expression data, including bulk and single-cell RNA sequencing data. A comprehensively integrated and well-curated erythropoiesis-specific database will greatly facilitate the mining of gene expression data and enable large-scale research of erythropoiesis and erythroid related diseases. Here, we present EryDB, an open-access and comprehensive database dedicated to the collection, integration, analysis, and visualization of transcriptomic data for erythropoiesis and erythroid-related diseases. Currently, the database includes expertly curated quality-assured data of 3,803 samples and 1,187,119 single cells derived from 107 public studies of three species (*Homo sapiens*, *Mus musculus*, and *Danio rerio*), nine tissue types, and five diseases. EryDB provides users with the ability to not only browse the molecular features of erythropoiesis between tissues and species, but also perform computational analyses of single-cell and bulk RNA sequencing data, thus serving as a convenient platform for customized queries and analyses. EryDB v1.0 is freely accessible at https://ngdc.cncb.ac.cn/EryDB/home.

## Introduction

Erythrocytes (red blood cells), the most abundant of the blood cells, are responsible for transporting oxygen from lungs to other tissues [1]. Erythrocyte impairment is attributed to acquired/hereditary erythropoietic diseases in which abnormalities occur in causative genes, including genes that encode erythrocyte membrane proteins, globins, and enzymes, as well as genes that are required for erythropoiesis [2]. Erythropoiesis is a finely regulated and complicated process that involves multiple transformations from hematopoietic stem cells to mature red blood cells at hematopoietic sites from the embryonic to adult stages [3]. Moreover, the current understanding of erythropoiesis and the pathophysiology of erythropoietic defects is limited. The underlying mechanisms need to be better defined to improve *in vitro* regeneration of red blood cells [4, 5] for the development of clinical treatments [3, 6]. Large amounts of multi-omics data are now available [7–13], providing unprecedented insights into erythroid biology and generating many novel hypotheses. However, most of the omics data are available as raw data deposited in public repositories, such as the Genome Sequence Archive (GSA) [14] and Sequence Read Archive (SRA) [15], making it challenging for clinicians and researchers to reanalyze the data and gain biological insights [16]. Although some research groups have made their erythroid profile data web-accessible [9, 11, 17], no erythropoiesis-specific database that can facilitate the reuse of omics data is available to the erythroid biology community. An authoritative and integrated omics database that contains the molecules, pathways, and events involved in erythropoiesis will help to mitigate these shortcomings and accelerate research in this area.

Here, we present EryDB (https://ngdc.cncb.ac.cn/EryDB/home), a database that contains curated, quality-assured, and pre-analyzed transcriptome data of erythropoiesis and erythroid-related diseases. The EryDB platform allows users to search, browse, and analyze gene expression profiles or related data among various tissues or experimental types. Furthermore, gene expression changes under various pathological conditions can be analyzed towards developing diagnostic and prognostic tools for various erythropoietic diseases.

In this paper, we describe the breadth of the transcriptomic data available in EryDB, which includes bulk and single-cell RNA sequencing (RNA-seq) data. The datasets are classified into four major erythropoiesis-related categories, namely erythroid differentiation, genes, compounds, and diseases that users can mine to obtain erythroid-specific information. Customized filtering of the data can be performed under any of the four categories. These tools allow users to easily retrieve erythroid-specific data of interest to them. Moreover, EryDB provides a functional module named the Erythroid Atlas that can be used to integrate and compare erythropoiesis-related datasets from different tissues, namely bone marrow, peripheral blood, cord blood, and embryos. The rapid data query and analysis capabilities provided by EryDB enable efficient data mining for basic and clinical research.

## Data collection and processing

### Data collection and classification

The process that was used to construct EryDB is outlined in **Figure 1**. Transcriptome data of erythroid cells were obtained from the Gene Expression Omnibus (GEO) [18], Sequence Read Archive (SRA) [15], and Genome Sequence Archive (GSA) [14] databases using the keywords “erythroid” or “erythrocyte”. We also conducted a PubMed search to identify recent single-cell transcriptome studies using the keywords ’peripheral blood mononuclear cell’ or ‘hematopoiesis’ and ’erythroid ’ as well as ’single cell’. Furthermore, we screened the titles, abstracts, and sample information to identify studies that specifically performed gene expression profiling of erythroid cells. For the single-cell RNA-seq (scRNA-seq) data, only erythroid-related studies with > 300 cells were included. We also included published [19–22] and unpublished data from our laboratory. Finally, we assigned a unique custom ID to each dataset in the Erythroid Database to facilitate user-friendly searches. Additionally, we retained the data source as part of the dataset metadata.

**Figure 1.**
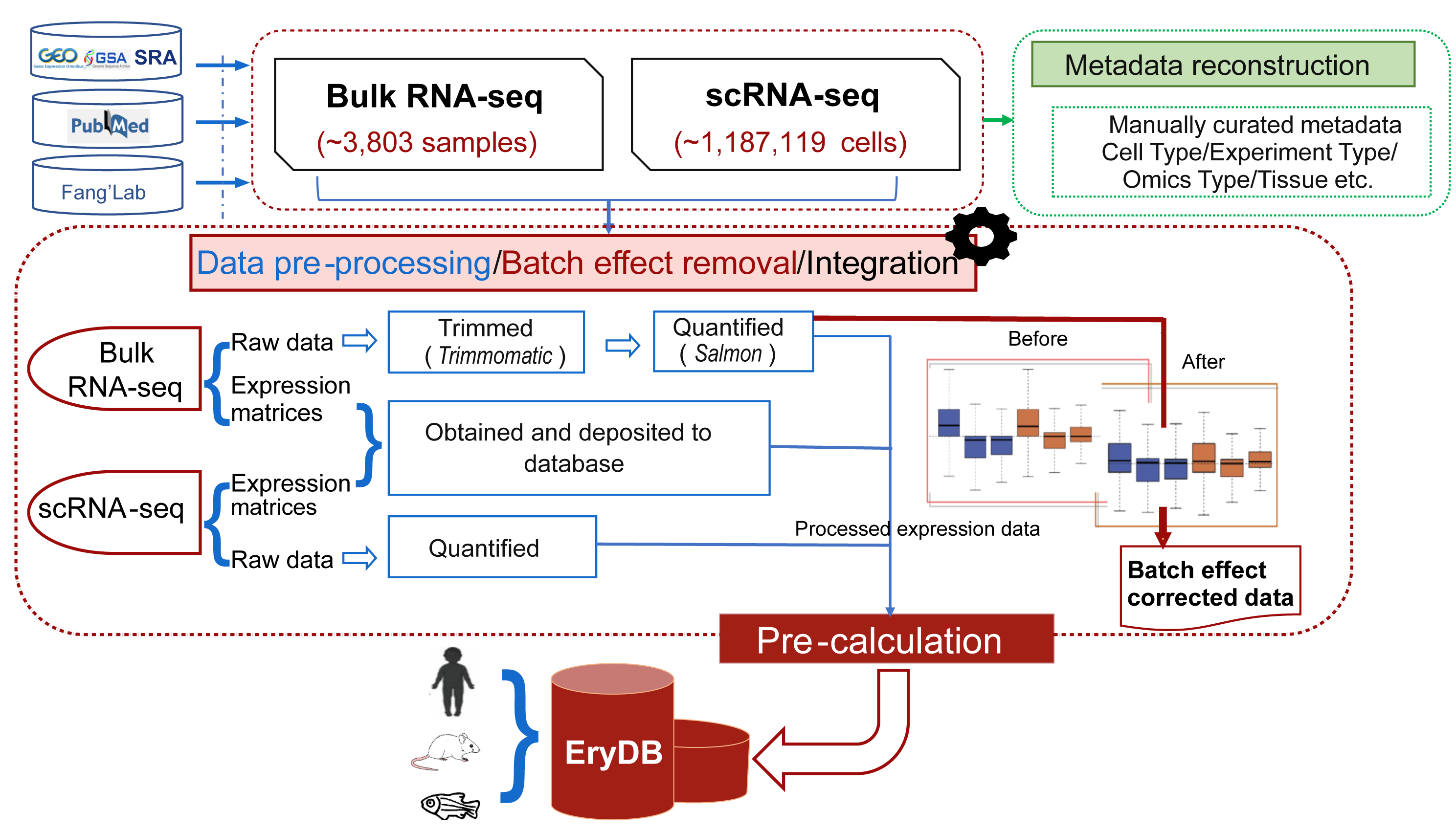
Construction process of EryDB. Transcriptome data for erythroid cells were obtained from GEO, Gene Expression Omnibus, SRA , Sequence Read Archive SRA, and GSA, Genome Sequence Archive. Metadata were downloaded from GEO, SRA, GSA and manually curated by experts in erythropoiesis.

### Metadata curation

Metadata were downloaded from GEO, SRA, GSA and manually curated by experts in erythropoiesis (Figure 1). Repetitive and redundant information was removed. Each dataset was manually annotated with biologically relevant information, including model organism, sample tissue, experiment type, and omics type. A description of each study, GEO or GSA identifiers, and references to the original publications were also included in the metadata. All the datasets are labeled with species, tissue, and experiment type (*in vivo*/*in vitro*).

### Pre-processing and quality control of bulk transcriptomic data

For the RNA-seq datasets, we obtained raw sequencing data whenever possible; in cases where raw sequencing data were unavailable, we included pre-processed data or raw count files. We employed an identical processing pipeline for datasets with access to raw sequencing data. The adaptor and low-quality sequences were removed using *Trimmomatic* (v.0.36) [23]. Gene expression was quantified using *Salmon* (v.1.4.0) [24] (Figure 1). Trimmed mean of M-values and Voom normalization were performed using the *EdgeR* (v.3.42.4) [25] and *limma* (v.3.48.3) [26] packages in R (v.4.3.0), respectively. For the integrated datasets within Erythroid Atlas in EryDB, all samples have been processed using the same analysis pipeline and then removed batch effects before proceeding to downstream analysis. Gene annotations for *Homo sapiens* (GRCh38), *Mus musculus* (GRCm38), and *Danio rerio* (Zv9) were sourced from Ensemble (release-91).

### Pre-processing and quality control of single-cell transcriptomic data

For the scRNA-seq dataset, we downloaded the processed expression matrices when raw sequencing data were unavailable. The raw sequencing data was preprocessed using the methods described in the original publication, including quality control and normalization. Low-quality cells (expressing < 300 genes) and mitochondrial genes ( > 20% of the reads mapped to mitochondrial genes) were removed. Library size normalization was performed using the *Seurat* (v.4.3.0) [27] R package. The batch effects of combined expression data were removed using *Seurat* CCA function or Harmony (v.0.1.1) [28]. For the integrated single-cell datasets within Erythroid Atlas in EryDB, all raw sequencing data be pre-processed by an identical processing pipeline and the comparative analyses were performed after integration and batch correction .

## Implementation

EryDB is an interactive web application built using the Python/*Flask* (v.1.0.2) web framework. We used *Vue*, *Plotly*, *Encharts*, and *Highcharts* with Python for data visualization, and *iView.Table* to construct searchable tables.

## Database content and usage

The EryDB database is a free public biological resource for erythropoiesis-related research. Users can browse datasets, examine the expression of erythropoiesis-related genes, and compare transcriptome differences and cell variations in the context of erythropoiesis and erythroid-related diseases.

### Data records

In the current release, EryDB contains high-quality analytical results of 118 datasets and 3,803 samples from 107 studies of three species, *H. sapiens*, *M. musculus*, and *D. rerio*. Fifteen of these studies were scRNA-seq studies involving 1,187,119 single cells; the others were bulk RNA-seq studies. Among the datasets, 60 (50.85%), 49 (41.53%), and 9 (7.63%) were for *H. sapiens, M. musculus*, and *D. rerio*, respectively. The 118 datasets were associated with 11 tissue types: bone marrow, cord blood, peripheral blood, spleen, embryo, fetal tissue, heart, kidney, induced pluripotent stem cells (iPSC), cell line, and other (unknown). Data statistics for EryDB are shown in **Table 1**. The accession numbers for all the erythroid-related datasets are listed in **Table S1**.

**Table 1.** Statistics of the contents of EryDB.

| Species | Tissue | Omics type | No. of studies | No. of datasets | No. of samples | No. of single cells |
| --- | --- | --- | --- | --- | --- | --- |
| *Homo sapiens* | Fetal tissue<br>(e.g., liver, spleen, kidney) | scRNA-seq | 3 | 3 | 98 | 434,156 |
|  |  | Bulk RNA-seq | 5 | 5 | 80 | / |
|  | Bone marrow | scRNA-seq | 6 | 7 | 130 | 267,566 |
|  |  | Bulk RNA-seq | 3 | 6 | 88 | / |
|  | Peripheral blood | scRNA-seq | 1 | 1 | 2 | 26,740 |
|  |  | Bulk RNA-seq | 10 | 7 | 474 | / |
|  | Induced pluripotent stem cells | scRNA-seq | 2 | 2 | 731 | 27,666 |
|  |  | Bulk RNA-seq | 1 | 1 | 5 | / |
|  | Cord blood | scRNA-seq | 2 | 2 | 7 | 29,920 |
|  |  | Bulk RNA-seq | 13 | 13 | 142 | / |
|  | Cell line | Bulk RNA-seq | 6 | 6 | 82 | / |
|  | Other (unknown) | Bulk RNA-seq | 1 | 1 | 6 | / |
| *Mus musculus* | Fetal tissue<br>(e.g., liver, yolk sac, caudal half) | scRNA-seq | 1 | 1 | 6 | 353,528 |
|  |  | Bulk RNA-seq | 18 | 22 | 542 | / |
|  | Bone marrow | scRNA-seq | 2 | 1 | 9 | 28,224 |
|  |  | Bulk RNA-seq | 10 | 15 | 1,162 | / |
|  | Cell line | Bulk RNA-seq | 4 | 6 | 46 | / |
|  | Spleen | Bulk RNA-seq | 4 | 6 | 43 | / |
|  | Embryo | Bulk RNA-seq | 3 | 3 | 24 | / |
|  | Heart | Bulk RNA-seq | 2 | 1 | 6 | / |
| *Danio rerio* | Fetal tissue | scRNA-seq | 1 | 1 | 1 | 19,319 |
|  | Embryo | Bulk RNA-seq | 7 | 6 | 93 | / |
|  | Kidney | Bulk RNA-seq | 2 | 2 | 24 | / |
| **Sum** |  |  | 107 | 118 | 3,803 | 1,187,119 |

### User-friendly search modules for data queries

EryDB provides three methods for datasets searches. First, a quick-search box on the home page that allows real-time queries by specifying genes, tissues, dataset ID, organism name or PubMed Unique Identifier (PMID). Second, category-specific searches by Erythroid Differentiation, Genes, Compounds, or Diseases that are also available on the home page: Erythroid Differentiation—the process of erythroid differentiation is subdivided into 12 cell types (**Figure 2A**), and datasets that contain a cell type of interest are listed by clicking of the cell type; Genes—important genes related to erythroid differentiation and development are summarized as a keyword cloud and a sorted table, and datasets associated with a gene of interest can be accessed by clicking on the gene name (**Figure 2B**); Compounds—molecular compounds related to erythroid differentiation and development are listed (**Figure 2C**), and datasets associated with a compound of interest can be accessed by clicking on the compound name; and Diseases—the five main diseases related to erythropoiesis are available in 14 datasets (**Figure 2D**), and datasets associated with a disease of interest can be accessed by clicking on the disease name. The query results from these four categories are presented as tables that contains the dataset ID, species, tissue, and experiment type (**Figure 2E**). The table heading in each column is a drop-down box that enables further filtering of the datasets according to specific conditions. Third, target datasets can be selected by clicking on “Search” at the top of the home page and choosing one or more of the options on the navigation bar on the left-hand side of the page, namely cell type, reported gene, compound type, disease type, species, tissue, experiment type, and omics type (**Figure 2F**). Detailed information about the content of the datasets that can be searched under these conditions is listed in **Table S2**. The results of such queries are listed in tables that are similar to those obtained by category-specific searches.

**Figure 2.**
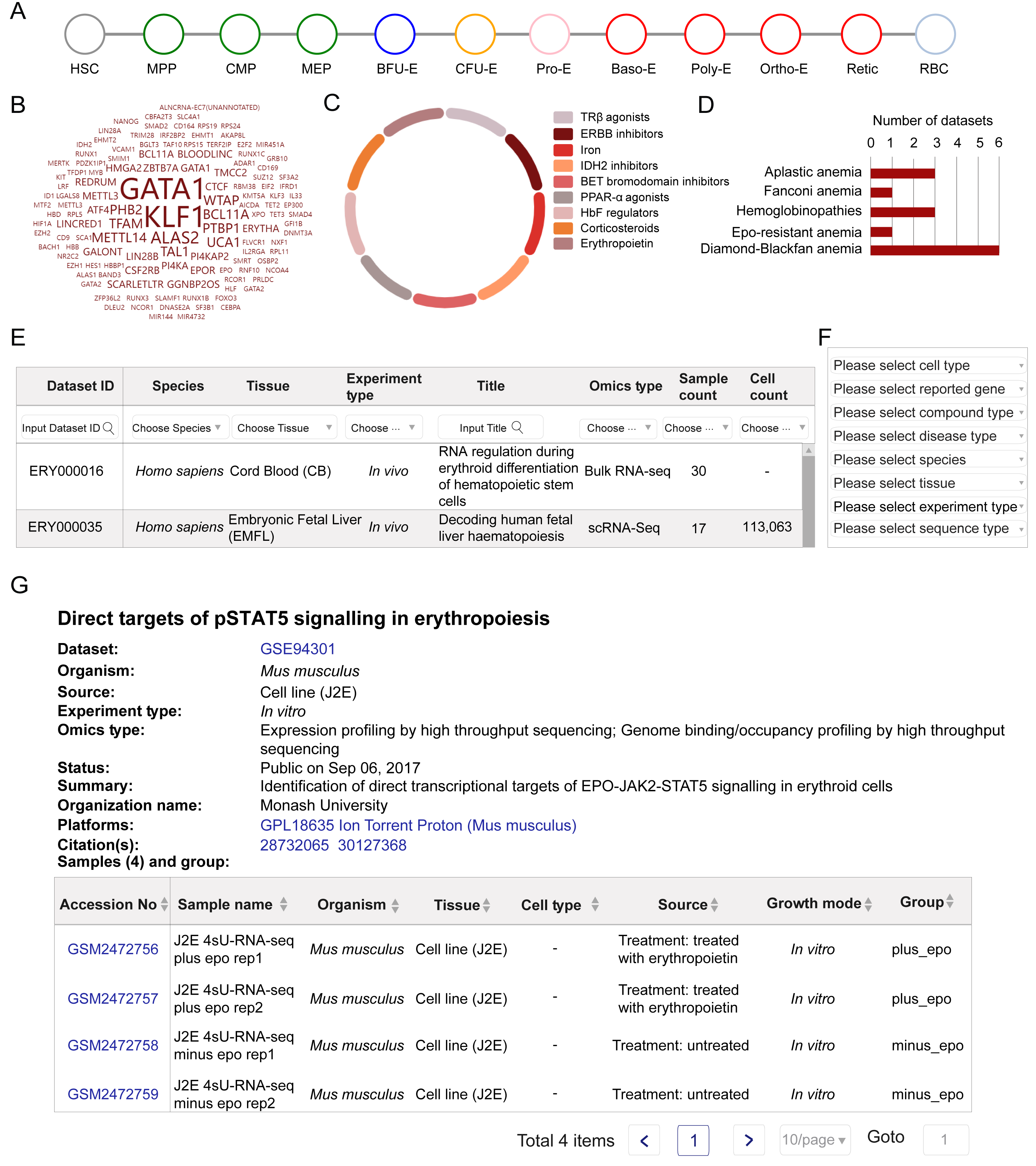
Dataset query strategies in EryDB. **A.** The Erythroid Differentiation category allows datasets to be filtered by cell type. **B.** The Genes category allows datasets to be filtered by selecting a gene of interest. **C.** The Compounds category allows datasets under the action of a certain molecular compound to be obtained. **D.** The Diseases category allows datasets to be obtained by selecting a specific disease. **E.** Query results are presented as a table. **F.** Navigation bar on the left-hand side of the “Search” page lists the conditions for selecting target datasets. **G.** Detailed meta information included in the dataset overview page.

Clicking on any word in the results tables from all three types of searches expands the overview and the gene expression profiles of the dataset. The overview includes a research summary, research organization, sequencing platform information, specific sample information, data processing method, overall design introduction, data contributors, and contact information for the dataset (**Figure 2G**). The sample accession numbers are linked to the database source of the original data. Because of the different technologies used for bulk RNA-seq and scRNA-seq, the gene expression profile datasets from the two technologies are presented separately.

### Comprehensive multidimensional online data exploration

#### Bulk RNA-seq data exploration

EryDB provides four analysis modules for the bulk transcriptome datasets on the overview page: Expression Profile—analysis of expression changes for a gene, sample group, and dataset specified by users (**Figure 3A**). The detailed results are shown when the sample bar is selected; Principal Component Analysis—a two-dimensional (2D) plot of the principal component analysis results is obtained, where individual sample’s group are color-coded (**Figure 3B**); Differential Analysis—a volcano plot of differentially expressed genes (DEGs) between two user-specified groups and a browsable table below the plot are obtained (**Figure 3C**); and Enrichment Analysis— enrichment analysis results of DEGs between user-specified groups, including Gene Ontology (GO) and Kyoto Encyclopedia of Genes and Genomes (KEGG) functional and pathway annotations (**Figure 3D**).

**Figure 3.**
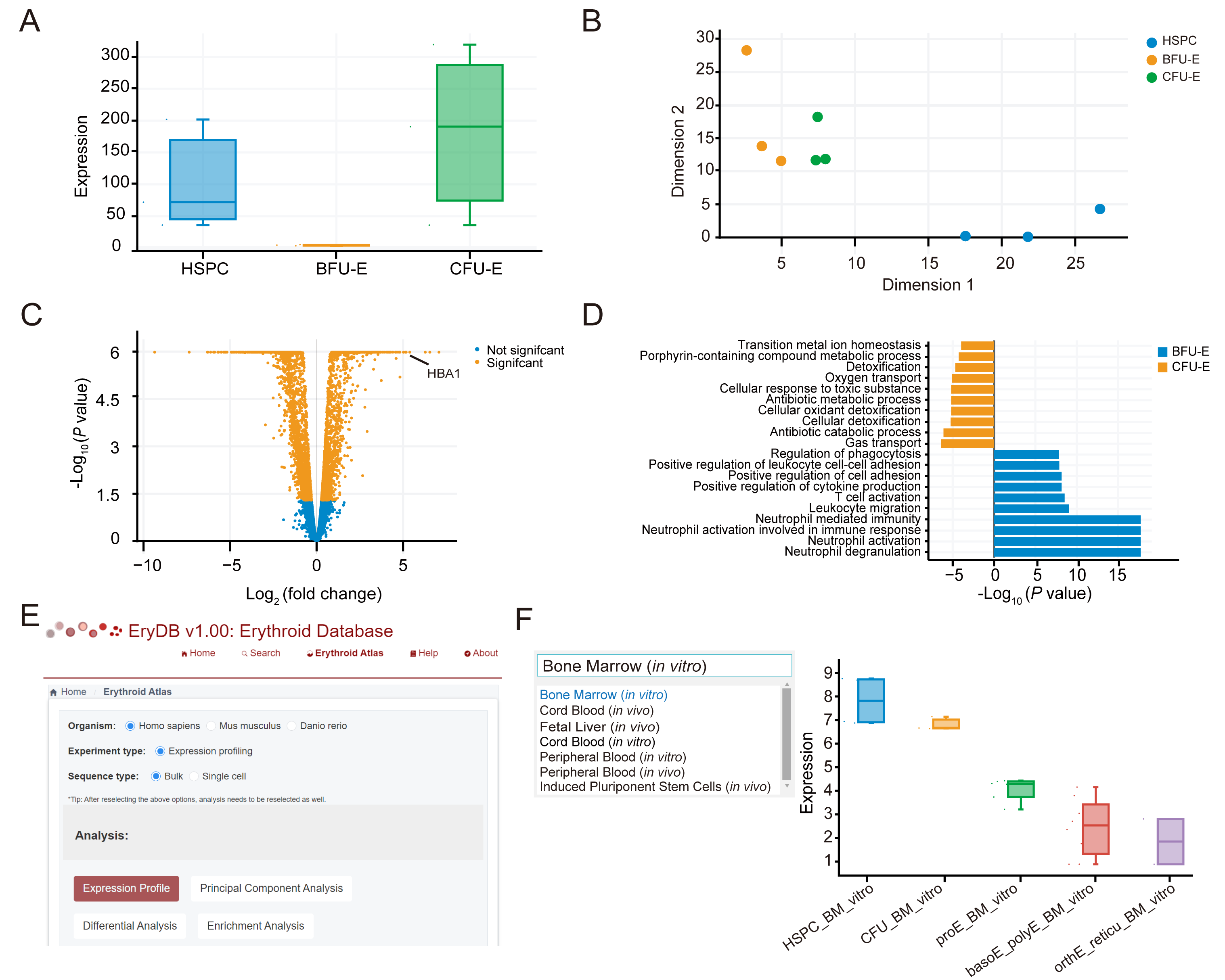
Statistical analysis function for bulk RNA-seq data. **A.** Expression profile of hemoglobin subunit gene *HBA1* in dataset GSE128268 as an example to demonstrate the functions of EryDB. **B.** Principal component analysis of samples in the GSE128268 dataset. **C.** Volcano plot of DEGs, differentially expressed genes between samples from the groups of CFU-E, the colony-forming unit erythroid and BFU-E, the burst forming unit-erythroid in GSE128268. **D.** Enrichment analysis (biological process terms of Gene Ontology) of DEGs between samples from the CFU-E and BFU-E groups in GSE128268. **E.** Screenshot of the Erythroid Atlas page. Datasets for different differentiation stages can be explored on this page. **F.** Expression of the transcription factor gene *RUNX2* among different developmental stages across multiple datasets of bone marrow tissue as an example. The tissue of interest can be selected from the drop-down box on the left-hand side.

To demonstrate the database functions, we applied EryDB to the GEO dataset GSE128268, which is a bulk RNA-seq dataset of an erythroid cell culture of human umbilical cord-derived CD34+ hematopoietic stem and progenitor cells [29]. We found that the hemoglobin subunit gene *HBA1* was highly expressed at the colony-forming unit erythroid (CFU-E) stage (Figure 3A). Furthermore, the burst forming unit-erythroid (BFU-E) and CFU-E cells had transcriptome profiles that were clearly distinct, and both profiles were distinct from those of umbilical cord-derived CD34+ hematopoietic stem and progenitor cells (Figure 3B). To determine which pathways were most upregulated in CFU-E cells, we compared the gene expression signatures between CFU-E and BFU-E cells and found that gas and oxygen transport were upregulated in CFU-E cells (Figure 3D). Analysis of the DEGs showed that hemoglobin markers (e.g., *HBA1*, *HBA2*, and *HBG2*) were among the top 25 most upregulated genes at the CFU-E stage (Figure 3C). Together, these results suggest that CFU-E cells undergo a switch that results in a hemoglobin-based oxygen-carrier phenotype.

The Erythroid Atlas page in EryDB allows comparisons to be made between transcriptome datasets associated with erythroid development. Users can explore the gene expression profiles of different differentiation systems in humans, mice, and zebrafish (**Figure 3E**). For example, changes in the expression of the transcription factor gene *RUNX2* at different erythroid differentiation stages in humans (*in vivo*) or changes induced *in vitro* can be browsed (**Figure 3F**). The results were obtained by the integration of 11 datasets that contain a total of 207 samples. All the data are analyzed and normalized in a standard pipeline to eliminate the batch effect of the data sources. Moreover, DEGs can be identified from volcano plots and tables between any two stages of the erythroid differentiation process by selecting corresponding tags in the drop-down box, which enables users to identify genes of interest for further study.

#### scRNA-seq data exploration

EryDB provides six analysis modules for scRNA-seq data: Visualization & Feature— enables dimension reduction analysis and signature expression queries among cells. Gene expression among single cells is displayed as interactive t-distributed stochastic neighbor embedding (t-SNE) or uniform maximal approximation projection (UMAP) plots (**Figure 4A** and **4B**), and the expression of specific genes among cells can be queried in an extra t-SNE plot (**Figure 4C**); Marker & Enrichment—provides marker gene expression of cell types as a heatmap (**Figure 4D**) and enables functional enrichment analysis of genes in cell populations. The KEGG pathway or GO enrichment results for each stage are presented as bar graphs (**Figure 4E**); Difference & Enrichment—the expression of DEGs in the same cell type from different group is presented in a volcano plot and the functional enrichment results of these DEGs are presented in a bar plot; Differentiation Trajectory—pseudotime analysis of cells where pseudotime trajectories of cell differentiation are visualized in 2D space (**Figure 4F**), and cell differentiation trajectories can be displayed by cell type (**Figure 4G**); Cell– Cell Interaction—the interaction of ligands and receptors between different cell types can be viewed as a circus plot, where the thickness of the lines indicates the interaction strength. The communication types include growth factors, cytokines, checkpoints, and others, which can be queried by selection in the box provided (**Figure 4H**); and Cell–Cell Communication—cellular communication is visualized as a river diagram in which each pattern represents different levels of communication (**Figure 4I**). User can view the interaction of a signaling pathway of interest (**Figure 4J**) and the contribution of specific ligand-receptor pairs in this pathway (**Figure 4K**). Together, these analyses cover most of the statistical bases of single-cell transcriptomes.

**Figure 4.**
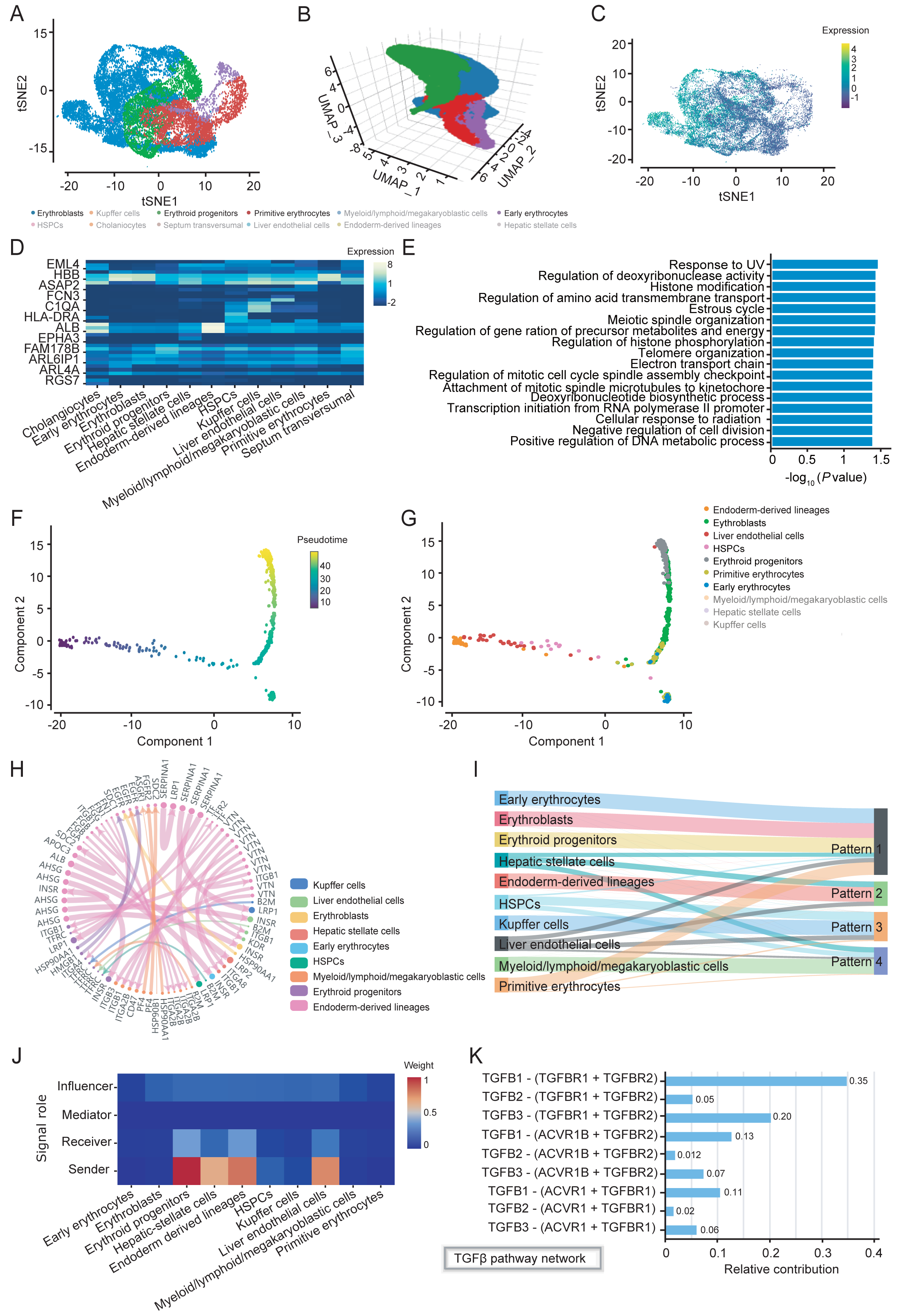
Statistical analysis function for scRNA-seq data. **A.** Interactive two-dimensional t-SNE, t-distributed stochastic neighbor embedding plot of the single cells in the W9 group in dataset CRA002443 as an example to demonstrate the functions of EryDB; cell clusters are color-coded. **B.** Interactive three-dimensional UMAP, uniform maximal approximation projection plot of the single cells in the W9 group; cell clusters are color-coded. **C.** Expression of the cyclin gene *CCNB1* among erythroblasts as an example. Panels A, B, and C illustrate the usage of the Visualization & Feature module. **D.** Expression of marker genes at different cell stages. **E.** GO, Gene Ontology and KEGG, Kyoto Encyclopedia of Genes and Genomes functional and pathway enrichment for genes in the erythroblasts cluster. The cluster and enriched type can be selected in the drop-down boxes in the Marker & Enrichment module. **F.** Pseudotime trajectories of cell differentiation visualized in 2D. **G.** Cell differentiation trajectories displayed by cell type. Panels F and G show the usage of the Differentiation Trajectory module. **H.** Interaction of ligands and receptors between different cell types. Interaction of the top 40 ligands and receptors pairs is shown. The communication type can be defined in the drop- down boxes in the Cell–Cell Interaction module. **I.** Cell–Cell communication is visualized as a river diagram. **J.** Interaction of the TGFβ signaling pathway between cells as an example. **K.** Contribution of specific ligand-receptor pairs in the TGFβ signaling pathway. Panels I, J, and K illustrate the usage of the Cell–Cell Communication module.

To demonstrate the database functions, we applied EryDB to the GSA dataset CRA002443, which is a single-cell transcriptome dataset of human 5–19 week post- coitum fetal tissue (W5–19) [7], and explored the erythroid lineage properties. A total of 12 cell clusters were identified in the W9 sample, and the number of erythroid cell lineages in this sample was higher than the numbers in the other samples (Figure 4A and 4B). Expression of the cyclin gene *CCNB1* was upregulated in erythroblasts (Figure 4C). Markers for each cell population are plotted as a heatmap (Figure 4D) that can be enlarged for convenient browsing of gene expression. The hemoglobin gene *HBB* was highly expressed in erythroid cell lineages in the W9 sample, and genes expressed in erythroblasts were enriched in several biological processes, including response to UV and regulation of deoxyribonuclease activity (Figure 4E). In the W9 group, endoderm-derived cell lineages had the strongest cell-cell interactions among the top 40 pairs of ligands and receptors, but they showed much less interactions in early erythroblasts (Figure 4H). Four communication patterns were identified among different cell types (Figure 4I). The transforming growth factor TGFβ signaling pathway in two of the patterns (patterns 1 and 2) was communicated between most of the cells by multiple ligands and receptors (Figure 4J and 4K).

Similarly, the Erythroid Atlas page provides integrative analysis via six modules across differentiated systems from the scRNA-seq datasets.

In summary, these examples demonstrate the power of EryDB to efficiently test novel hypotheses using publicly available data of genes that regulate erythroid differentiation and development. All visualizations can be downloaded as SVG or PNG files, and related result data tables can be downloaded in comma-separated value (CSV) format or Excel sheets. For advanced users, the integrated expression matrix and metadata can be downloaded for personalized analysis. Besides, the analysis results can be filtered with adjustable parameters. The fully functional design of EryDB is intended to facilitate the widespread reuse of publicly available datasets to address novel and unresolved questions and allow comparative analyses of similar studies to resolve study-specific biases.

## Discussion

A survey of erythroid-related omics databases confirmed that the currently available databases are important resources that can be used to interpret gene regulation or mutations related to erythropoiesis and erythroid-related diseases. Some erythroid- related databases are no longer accessible (e.g., EpoDB [30], Hembase [31], and ErythronDB [32]). Accessible databases (e.g., dbRBC [33], RBCmembrane [34], and BloodSpot [35, 36]) that contain specific omics data associated with red blood cells (RBCs) (**Table S3**) tend to lack diversity in their data sources, species covered, or functional analysis tools, and do not contain single-cell sequencing data.

EryDB was designed to integrate most of the accessible public transcriptome datasets related to erythropoiesis and enable the investigation of gene expression dynamics in erythroid differentiation and erythropoiesis-related diseases. Compared with the other RBC-related databases, EryDB has more user-friendly data search options, more extensive data sources, and more comprehensive analysis capabilities. In the sub- module search of the datasets, users can select the search that meets their research interests by selecting the relevant module. For example, in the Disease module, researchers can easily examine the gene expression changes under certain pathological conditions for blood disease diagnosis [37–40] and in the Compounds module, users can obtain information about the small molecules that promote or inhibit erythroid differentiation. The *in vitro* RBC regeneration data from our laboratory are unique in EryDB and can be integrated with *in vivo*-derived erythroid cell data. This allows users to compare the differences and mechanisms underlying erythropoiesis from different global sources. EryDB also provides comprehensive functional analysis tools. For example, cell-cell interaction analysis based on scRNA-seq datasets generated a broad spectrum of information on the interactions between erythroid cells and the other cell types. We found that *in vivo* erythroid cells had stronger cellular interactions with the other cell types than *in vitro* erythroid cells had.

EryDB will be useful not only for computational biologists but also for bench clinicians interested in erythropoietic disorders and for researchers involved in basic erythropoiesis-related research. In future releases of EryDB, the transcriptomic data will be continuously updated, and other types of omics data will be added. We will also develop new analytical feature for further exploration of the multi-omics data.

## Data availability

EryDB v1.0 is freely accessible at https://ngdc.cncb.ac.cn/EryDB/home.

## CRediT authorship statement

**Guangmin Zheng:** Software, Visualization, Writing - original draft, Writing - review & editing. **Song Wu**: Software, Visualization, Validation, Writing - review & editing. **Zhaojun Zhang:** Data curation, Resources, Investigation, Validation, Writing - review & editing. **Zijuan Xin:** Data curation, Resources, Investigation**. Lijuan Zhang:** Data curation, Resources, Investigation**. Siqi Zhao:** Validation. **Jing Wu:** Validation. **Yanxia Liu:** Validation. **Meng Li:** Validation. **Xiuyan Ruan:** Validation. **Yiming Bao:** Supervision, Project administration, Funding acquisition, Resources, Conceptualization. **Hongzhu Qu**: Validation, Writing - review & editing. **Xiangdong**

**Fang:** Supervision, Project administration, Funding acquisition, Resources, Conceptualization. All authors reviewed and approved the final manuscript.

## Acknowledgements

We would like to thank Yunxiao Ren, Rudan Xiao for their assistance in RNA data curation. This research was supported by the Strategic Priority Research Program of the Chinese Academy of Sciences (Grant No. XDA16010602), National Natural Science Foundation of China (Grant Nos. 81870097, 82070114, 82270126), the National Key Research and Development Program of China (Grant No. 2022YFC2406803). The authors would like to thank Liwen (www.liwenbianji.cn) and Editage (www.editage.cn) for the English language review.

---

**Table S1 All erythroid-related datasets in EryDB**

**Table S2 Content of the datasets that can be searched by users**

**Table S3 Other publicly available erythropoiesis-related databases**

## References

[1] Palis J. Primitive and definitive erythropoiesis in mammals. Front Physiol 2014; 5:3.

[2] Di Pierro E, Brancaleoni V, Granata F. Advances in understanding the pathogenesis of congenital erythropoietic porphyria. Br J Haematol 2016; 173(3):365–379.

[3] Caulier AL, Sankaran VG. Molecular and cellular mechanisms that regulate human erythropoiesis. Blood 2022; 139(16):2450–2459.

[4] Batta K, Menegatti S, Garcia-Alegria E, Florkowska M, Lacaud G, Kouskoff V. Concise Review: Recent Advances in the In Vitro Derivation of Blood Cell Populations. Stem Cells Transl Med 2016; 5(10):1330–1337.

[5] Christaki EE, Politou M, Antonelou M, Athanasopoulos A, Simantirakis E, Seghatchian J et al. Ex vivo generation of transfusable red blood cells from various stem cell sources: A concise revisit of where we are now. Transfus Apher Sci 2019; 58(1):108–112.

[6] Weiss MJ, dos Santos CO. Chaperoning erythropoiesis. Blood 2009; 113(10):2136–2144.

[7] Wang X, Yang L, Wang YC, Xu ZR, Feng Y, Zhang J et al. Comparative analysis of cell lineage differentiation during hepatogenesis in humans and mice at the single-cell transcriptome level. Cell Res 2020; 30(12):1109–1126.

[8] Yang D, Jang I, Choi J, Kim MS, Lee AJ, Kim H et al. 3DIV: A 3D-genome Interaction Viewer and database. Nucleic Acids Res 2018; 46(D1):D52–D57.

[9] Tusi BK, Wolock SL, Weinreb C, Hwang Y, Hidalgo D, Zilionis R et al. Population snapshots predict early haematopoietic and erythroid hierarchies. Nature 2018; 555(7694):54-60.

[10] Weinreb C, Wolock S, Tusi BK, Socolovsky M, Klein AM. Fundamental limits on dynamic inference from single-cell snapshots. Proc Natl Acad Sci U S A 2018; 115(10):E2467–E2476.

[11] Popescu DM, Botting RA, Stephenson E, Green K, Webb S, Jardine L et al. Decoding human fetal liver haematopoiesis. Nature 2019; 574(7778):365-371.

[12] Cao J, O’Day DR, Pliner HA, Kingsley PD, Deng M, Daza RM et al. A human cell atlas of fetal gene expression. Science 2020; 370(6518).

[13] Vanuytsel K, Matte T, Leung A, Naing ZH, Morrison T, Chui DHK et al. Induced pluripotent stem cell-based mapping of beta-globin expression throughout human erythropoietic development. Blood Adv 2018; 2(15):1998–2011.

[14] Members C-N, Partners. Database Resources of the National Genomics Data Center, China National Center for Bioinformation in 2022. Nucleic Acids Res 2022; 50(D1):D27-D38.

[15] Leinonen R, Sugawara H, Shumway M, International Nucleotide Sequence Database C. The sequence read archive. Nucleic Acids Res 2011; 39(Database issue):D19-21.

[16] Rung J, Brazma A. Reuse of public genome-wide gene expression data. Nat Rev Genet 2013; 14(2):89–99.

[17] Reyes M, Filbin MR, Bhattacharyya RP, Billman K, Eisenhaure T, Hung DT et al. An immune-cell signature of bacterial sepsis. Nat Med 2020; 26(3):333–340.

[18] Barrett T, Troup DB, Wilhite SE, Ledoux P, Rudnev D, Evangelista C et al. NCBI GEO: mining tens of millions of expression profiles--database and tools update. Nucleic Acids Res 2007; 35(Database issue):D760-765.

[19] Xin Z, Zhang W, Gong S, Zhu J, Li Y, Zhang Z et al. Mapping Human Pluripotent Stem Cell-derived Erythroid Differentiation by Single-cell Transcriptome Analysis. Genomics Proteomics Bioinformatics 2021; 19(3):358–376.

[20] Ren Y, Zhu J, Han Y, Li P, Wu J, Qu H et al. Regulatory association of long noncoding RNAs and chromatin accessibility facilitates erythroid differentiation. Blood Adv 2021; 5(23):5396–5409.

[21] Ding N, Xi J, Li Y, Xie X, Shi J, Zhang Z et al. Global transcriptome analysis for identification of interactions between coding and noncoding RNAs during human erythroid differentiation. Front Med 2016; 10(3):297–310.

[22] Yang Y, Wang H, Chang KH, Qu H, Zhang Z, Xiong Q et al. Transcriptome dynamics during human erythroid differentiation and development. Genomics 2013; 102(5-6):431–441.

[23] Bolger AM, Lohse M, Usadel B. Trimmomatic: a flexible trimmer for Illumina sequence data. Bioinformatics 2014; 30(15):2114–2120.

[24] Patro R, Duggal G, Love MI, Irizarry RA, Kingsford C. Salmon provides fast and bias-aware quantification of transcript expression. Nat Methods 2017; 14(4):417–419.

[25] Robinson MD, McCarthy DJ, Smyth GK. edgeR: a Bioconductor package for differential expression analysis of digital gene expression data. Bioinformatics 2010; 26(1):139–140.

[26] Ritchie ME, Phipson B, Wu D, Hu Y, Law CW, Shi W et al. limma powers differential expression analyses for RNA-sequencing and microarray studies. Nucleic Acids Res 2015; 43(7):e47.

[27] Stuart T, Butler A, Hoffman P, Hafemeister C, Papalexi E, Mauck WM, 3rd et al. Comprehensive Integration of Single-Cell Data. Cell 2019; 177(7):1888–1902 e1821.

[28] Korsunsky I, Millard N, Fan J, Slowikowski K, Zhang F, Wei K et al. Fast, sensitive and accurate integration of single-cell data with Harmony. Nat Methods 2019; 16(12):1289–1296.

[29] Schulz VP, Yan H, Lezon-Geyda K, An X, Hale J, Hillyer CD et al. A Unique Epigenomic Landscape Defines Human Erythropoiesis. Cell Rep 2019; 28(11):2996–3009 e2997.

[30] Stoeckert CJ, Jr., Salas F, Brunk B, Overton GC. EpoDB: a prototype database for the analysis of genes expressed during vertebrate erythropoiesis. Nucleic Acids Res 1999; 27(1):200–203.

[31] Goh SH, Lee YT, Bouffard GG, Miller JL. Hembase: browser and genome portal for hematology and erythroid biology. Nucleic Acids Res 2004; 32(Database issue):D572-574.

[32] Kingsley PD, Greenfest-Allen E, Frame JM, Bushnell TP, Malik J, McGrath KE et al. Ontogeny of erythroid gene expression. Blood 2013; 121(6):e5–e13.

[33] Patnaik SK, Helmberg W, Blumenfeld OO. BGMUT: NCBI dbRBC database of allelic variations of genes encoding antigens of blood group systems. Nucleic Acids Res 2012; 40(Database issue):D1023-1029.

[34] An X, Mohandas N. Disorders of red cell membrane. Br J Haematol 2008; 141(3):367–375.

[35] Bagger FO, Sasivarevic D, Sohi SH, Laursen LG, Pundhir S, Sønderby CK et al. BloodSpot: a database of gene expression profiles and transcriptional programs for healthy and malignant haematopoiesis. Nucleic Acids Research 2016; 44(D1):D917–D924.

[36] Bagger FO, Kinalis S, Rapin N. BloodSpot: a database of healthy and malignant haematopoiesis updated with purified and single cell mRNA sequencing profiles. Nucleic Acids Research 2019; 47(D1):D881–D885.

[37] Masuda T, Wang X, Maeda M, Canver MC, Sher F, Funnell AP et al. Transcription factors LRF and BCL11A independently repress expression of fetal hemoglobin. Science 2016; 351(6270):285-289.

[38] Lee HY, Gao X, Barrasa MI, Li H, Elmes RR, Peters LL et al. PPAR-alpha and glucocorticoid receptor synergize to promote erythroid progenitor self-renewal. Nature 2015; 522(7557):474-477.

[39] Wan Y, Zhang Q, Zhang Z, Song B, Wang X, Zhang Y et al. Transcriptome analysis reveals a ribosome constituents disorder involved in the RPL5 downregulated zebrafish model of Diamond-Blackfan anemia. BMC Med Genomics 2016; 9:13.

[40] Deena Iskander GW, Guanlin Wang, Guanlin Wang. Single-cell profiling of human bone marrow progenitors reveals mechanisms of failing erythropoiesis in Diamond-Blackfan anemia. Science translational medicine 2021; 13(610):eabf0113.

