## Supplementary material for "EryDB: a transcriptomic profile database for erythropoiesis and erythroid-related diseases": Table S2

| **Table S2 Content of the datasets that can be searched by users** | | |
| --- | --- | --- |
| **Item** | **Amount** | **Content** |
| Cell types | 12 | Hematopoietic stem cell (HSC), multipotent progenitor cell (MPP), common myeloid progenitor (CMP), megakaryocyte-erythroid progenitor (MEP), burst-forming unit-erythroid (BFU-E), colony-forming unit-erythroid (CFU-E), proerythroblast (Pro-E), basophilic erythroblast (Baso-E), polychromatophilic erythroblast (Poly-E), orthochromatic erythroblast (Ortho-E), reticulocyte (Retic), and red blood cell (RBC) |
| Disease types | 5 | Fanconi anemia, diamond-blackfan anemia, hemoglobinopathies (including sickle cell disease and thalassemia), aplastic anemia, and epo-resistant anemia |
| Compound types | 9 | Thyroid hormone receptor (TRβ) agonists, epidermal growth factor receptor (ERBB) inhibitors, isocitrate dehydrogenase 2 (IDH2) mutant-specific inhibitors, peroxisome proliferator-activated receptor α (PPAR-α) agonists, fetal hemoglobin (HbF) regulators, BET bromodomain inhibitors, erythropoietin, iron, and corticosteroids |
| Species | 3 | *Homo sapiens*, *Mus musculus*, and *Danio rerio* |
| Tissue types | 11 | Bone marrow, cord blood, peripheral blood, spleen, embryo, fetal tissue, heart, kidney, induced pluripotent stem cells (iPSC), cell line, and other (unknown). |
| Experimental types | 2 | *In vivo* and *in vitro* |
| Omics techniques | 2 | Bulk and single-cell RNA sequencing (RNA-seq) |
