## Supplementary material for "EryDB: a transcriptomic profile database for erythropoiesis and erythroid-related diseases": Table S3

**Table S3 Other publicly available erythropoiesis-related databases**

| Name  (Year published) | Weblink | Species | Data type | Data  number | Main features |
| --- | --- | --- | --- | --- | --- |
| EpoDB  (1999) | [http://cbil.](http://cbil.humgen.upenn.edu/epodb/)  [humgen.](http://cbil.humgen.upenn.edu/epodb/)  [upenn.edu/](http://cbil.humgen.upenn.edu/epodb/)  [epodb/](http://cbil.humgen.upenn.edu/epodb/)  (no longer accessible) | Vertebrate | Sequence of genes | 185 entries with a complete gene sequence annotation | Database of genes expressed in vertebrate  red blood cells. |
| Hembase  (2004) | [http://hembase.](http://hembase.niddk.nih.gov/)  [niddk.nih.gov](http://hembase.niddk.nih.gov/)  (no longer accessible) | *Homo*  *sapiens* | Expressed sequence tags and referenced genes | 15,752 erythroblast expressed sequence tags and 380 referenced genes relevant | Integrated browser and genome portal designed for web-based examination of the human erythroid transcriptome. |
| ErythronDB  (2013) | [https://www.](https://www.cbil.upenn.edu/ErythronDB/)  [cbil.upenn.edu/](https://www.cbil.upenn.edu/ErythronDB/)  [ErythronDB/](https://www.cbil.upenn.edu/ErythronDB/)  (no longer accessible) | *Mus*  *musculus* | Gene expression (microarray) | 60 samples | Expression data from embryonic, fetal, and adult mouse. |
| dbRBC  (2012) | [https://www.](https://www.ncbi.nlm.nih.gov/gv/rbc/)  [ncbi.nlm.](https://www.ncbi.nlm.nih.gov/gv/rbc/)  [nih.gov/gv/rbc/](https://www.ncbi.nlm.nih.gov/gv/rbc/) | *Homo*  *sapiens* | DNA and clinical data | 1,251 alleles of all 40 gene loci | dbRBC and The Blood Group Antigen Gene Mutation Database (BGMUT) are publicly accessible platforms for DNA sequences and a curated set of blood mutation information. |
| RBC membrane  (2008) | [https://research.](https://research.nhgri.nih.gov/RBCmembrane/)  [nhgri.nih.gov/](https://research.nhgri.nih.gov/RBCmembrane/)  [RBCmembrane/](https://research.nhgri.nih.gov/RBCmembrane/) | *Homo*  *sapiens* | Mutation information of related genes | 5 genes with 173 mutation entries | Grouping of all mutation genes occurring in single or more kinds of inherited disorders of the erythrocyte membrane; associated with hemolytic anemia. |
| Blood Spot  (2019) | [https://servers.](https://servers.binf.ku.dk/bloodspot/)  [binf.ku.dk/](https://servers.binf.ku.dk/bloodspot/)  [bloodspot/](https://servers.binf.ku.dk/bloodspot/) | *Homo*  *sapiens* and *Mus musculus* | Bulk RNA-seq and scRNA-seq | >25,000 samples | Provides gene expression of healthy and malignant hematopoiesis in humans or mice; lacks erythroid omics datasets from different sources. |
